# Diverse Calcium Signaling in Astrocytes: Insights from a Computational Model

**DOI:** 10.1101/2024.07.03.601899

**Authors:** Thiago Ohno Bezerra, Antonio C. Roque

## Abstract

Astrocytes are complex cells that influence a variety of brain functions and behaviors. They are active cells that show a sharp increase in intracellular Ca^2+^ concentration in response to neurotransmitters (events called Ca^2+^ signals). The main source of intracellular Ca^2+^ is the stores in endoplasmic reticulum (ER), released by the activation of IP3 receptor channels on the ER membrane. As neurons, astrocytes from different brain regions show distinct Ca^2+^ signals. In addition, astrocytes can also show different patterns of Ca^2+^ responses. It is not yet clear how the diversity of astrocyte response emerge from the same mechanisms. Here we present a two variable astrocyte compartmental model for the Ca^2+^ and IP3 dynamics. We show that Ca^2+^ signals with different characteristics can emerge from changing the parameters associated with the Ca^2+^ and IP3 dynamics and the transmembrane current. We also show that global Ca^2+^ signals are required for the model to trigger different patterns of Ca^2+^ responses. The model present here can be used to simulate astrocytes from different brain regions and with distinct types of response.

## 1 Introduction

Astrocytes, one of the cell types in the nervous system, are involved several cognitive and brain functions, such as working memory [1–3], goal directed behaviors [3, 4], control of motor and social behaviors [5, 6], regulation of synaptic plasticity [2, 7] and neural synchronization [8, 9]. As neuron, astrocytes are also active cell, responding to glutamate, dopamine, and noradrenaline released by neurons [8, 10–14]. Activation of metabotropic receptors expressed on astrocyte by these neurotransmitters increases the astrocyte intracellular Ca^2+^ concentration and induce a Ca^2+^ spike-like responses [15, 16]. These responses are often called Ca^2+^ signals. During a Ca^2+^ signal, astrocytes release neuroactive molecules, called gliotransmitters, like glutamate and ATP [7, 17, 18] that modulate neuronal and synaptic activities [7, 8]. Astrocytes from different brain regions also show diversity in gene expression, morphology, and characteristics of the Ca^2+^ signals triggered [19–21]. For example, hippocampal astrocytes have a higher frequency of spontaneous Ca^2+^ than astrocytes in the striatum [19], and Ca^2+^ signals triggered in cortical astrocytes were longer compared to the response of hippocampal astrocytes [20].

The Ca^2+^ response of astrocytes is a complex phenomenon. The canonical mechanism generating the Ca^2+^ responses involve the release of Ca^2+^ from internal stores in the mitochondria and in the endoplasmic reticulum (ER) [10, 22]. IP_3_ channel receptors present in the ER membrane promote the efflux of Ca^2+^ to the cytosol [10, 22], increasing its intracellular concentration. Another mechanism for the generation of Ca^2+^ in astrocytes involves the transporter present in their membrane, such as the glutamate transporter, the Na^+^/K^+^-ATPase and the Ca^2+^/Na^+^-exchanger (NCX) [10, 23, 24]. The dominant mechanism generating the Ca^2+^ response in different regions of the same astrocyte depends on the surface-to-volume ratio. Regions with high surface-to-volume ratio also have ER with higher volume [25]. In these regions, release of internal Ca^2+^ from the ER stores contributes to the astrocytic response [24, 26]. In contrast, in regions with a low surface-to-volume ratio, the Ca^2+^ entry by the transporters and exchangers expressed on the astrocyte membrane has a stronger influence over the Ca^2+^ response [24, 26]. In this regard, Bindocci *et al*. described two types of astrocyte response, a local, spatially restricted response, and a global one, recruiting simultaneously several astrocyte regions [16]. So, the astrocyte morphology also affects the Ca^2+^ response.

Some reports showed that astrocytes trigger different types of Ca^2+^ signals [27–30]. In particular, a study combining experimental and modeling approaches, Taheri *et al*. [28] described four categories of Ca^2+^ signals: single peak, multipeak, plateau and long-lasting responses. By simulating a model for Ca^2+^ and IP_3_ dynamics, they showed different IP_3_ time courses led to distinct astrocytic Ca^2+^ responses. IP_3_ pulses that last longer produce the plateau response, while fast IP_3_ spikes are associated with single peak responses. However, it is also not yet clear how different IP_3_ time courses and Ca^2+^ signals can emerge from the same mechanisms. Since glutamate and neuromodulators have different release patterns and in a previous study of our group we showed that these neurotransmitters affect the astrocytic response in distinct ways [26], it is possible that the Ca^2+^ response type is controlled by the presence of neuromodulators. So, taking together, it is possible that both neurotransmitter type and morphological parameters affect the astrocyte activity and determine the type of Ca^2+^ responses. However, it is difficult to control experimentally both the site of stimulation and study the impact of astrocyte morphology.

The above mentioned difficulties can be avoided with mathematical models of astrocyte Ca^2+^ and IP_3_ dynamics. Most computational models of astrocytes are complex systems composed of several variables and parameters. In general, they are of the Li-Rinzel type model or the Höfer types models [31]. In the first case, the models account for the generation of Ca^2+^ signals considering the release of Ca^2+^ from intra-ER stores by the activation of IP_3_ channels. On the other hand, although simulating the same ER mechanism, the Höfer type models are reaction-diffusion models, detailing how spatial variables can also influence astrocytic Ca^2+^ response. Further studies also considered the transporter and Ca^2+^ channels [24, 28] and implemented different types of neurotransmitter receptors in the astrocyte model [15, 26, 32]. However, these models have several variables and parameters, making it difficult to study the astrocyte response diversity, use of dynamical system analysis to understand the model behavior and have a high the computational cost. Therefore, it is important to develop simplified models that capture the main characteristics of astrocytes and allow the analytical analysis of its dynamics.

Here we present a simplified astrocyte model composed of two variables that describe the intracellular Ca^2+^ and IP_3_ dynamics. The model developed here is consistent with both experimental data and other astrocyte models. The model triggers Ca^2+^ signals with different amplitude, duration and ISI by changing the Ca^2+^ and IP_3_ dynamics, and transmembrane current parameters. We show that glutamate, dopamine and astrocyte morphology can determine the type of Ca^2+^ signal triggered by the astrocyte.

## 2 Results

### 2.1 Characterization of the Simplified Model Response

To understand how the diversity of astrocytic response could emerge [28], we developed a simplified astrocyte model composed by both the ER- and the transporter-dependent mechanisms responsible for the Ca^2+^ response. The astrocyte model is a compartmental model and a bidimensional system in which the intracellular Ca^2+^ is described by a variable *c* (equation (9)) and IP_3_ concentration by a variable *i* (equation (15), Fig. 1a). We implemented the metabotropic glutamate receptor and the *α*_1_/dopaminergic receptors present in astrocyte membrane [13, 33] (equations (16) and (17)). The activation of these receptors promote the synthesis of IP_3_ (increase of *i* variable). Both glutamate and dopamine were simulated as Poisson processes with fixed frequency given in Hz (Extended Data Fig. and equations (21) and (22)). Ca^2+^ can also entry into the astrocyte cytosol through the Na^+^/Ca^2+^-exchanger (NCX, Fig. 1a) [10, 23, 24]. In the present model the ER volumes changes according to the compartment radius (Fig. 1b, equation (14)) controlling the strength of the ER- and NCX-dependent mechanisms. Here we used a linear morphological model (Fig. 1a) with the compartments connected by diffusion of Ca^2+^ and IP_3_ (equation (23)). The model full description and derivation can be found in the Model Description Section 3.

**Fig. 1.**
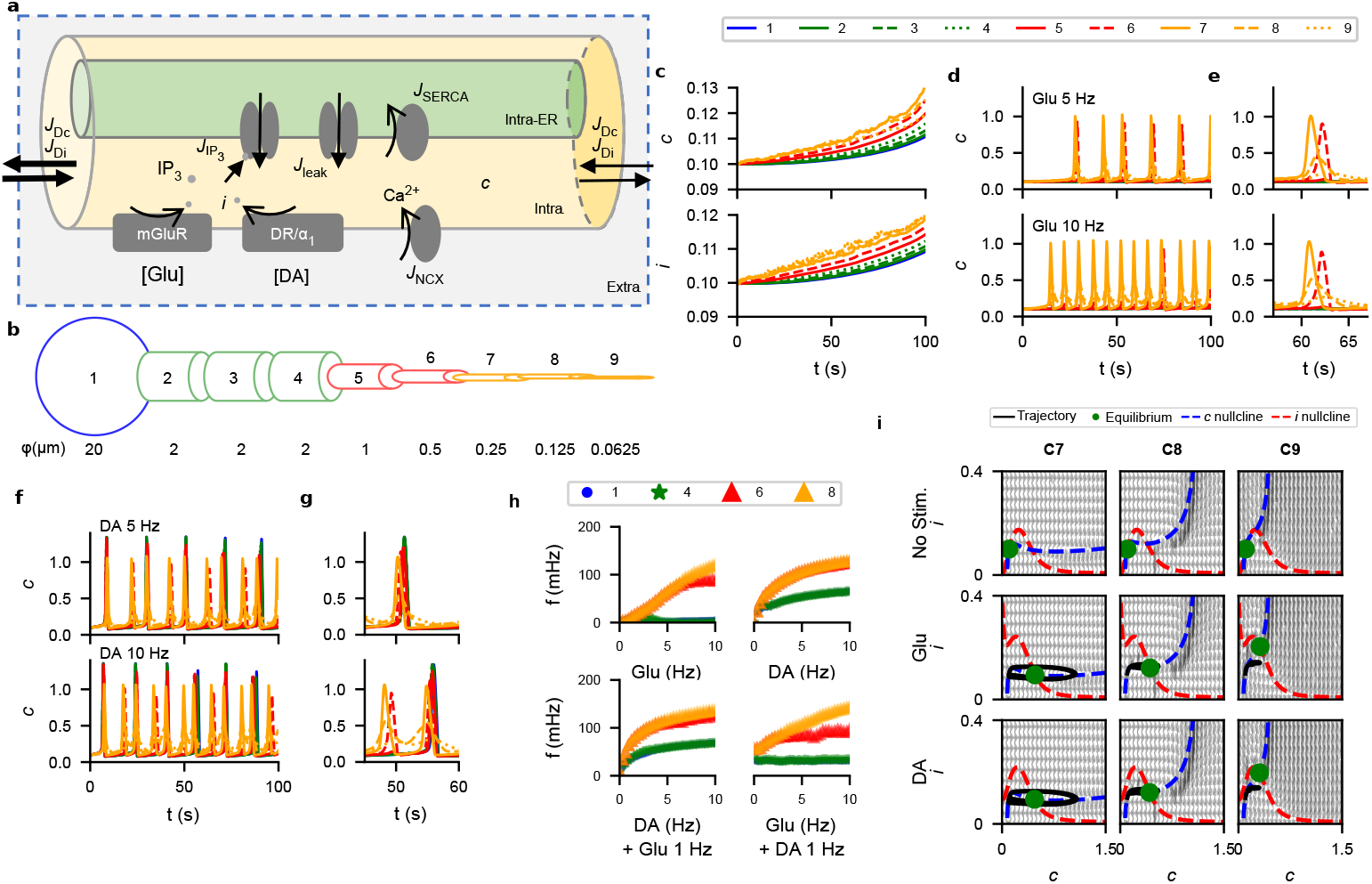
The simplified astrocyte model replicates the main characteristics of the Ca^2+^ responses in astrocytes. **a** schematic representation of a compartment of the astrocyte model with all variables and currents implemented. **b** representation of the linear morphology model. **c** intracellular Ca^2+^ and IP_3_ concentrations traces simulated with a glutamatergic input with 1 Hz. **d** model response to glutamatergic stimulation with 5 Hz (upper panel) or 10 Hz (lower panel). **e** amplification of a Ca^2+^ signal triggered by the glutamatergic input. **f, g** same as **d** and **e** for dopaminergic stimulation with 5 or 10 Hz. **h** input-response frequency curves for glutamatergic, dopaminergic or both stimuli. **i** model phase portrait of compartments 7, 8 and 9 with no stimulus, glutamate or dopamine. The phase portrait is plotted with the disconnect model.

To validate the model response, we first stimulated the model with a glutamate at the distal compartments, simulating synaptic inputs [34–36], and all compartments with dopamine, simulating its volume transmission [37]. A low frequency glutamatergic input (1 Hz) promoted the increase in the *i* variable, representing the synthesis of IP_3_ (Fig. 1c). This also increased the intracellular Ca^2+^ concentration, represented by the *c* variable. This effect is seem also in the non-stimulated compartments, showing that both Ca^2+^ and IP_3_ diffuse from the stimulated region to neighboring compartments. In contrast, glutamatergic inputs with 5 Hz and 10 Hz triggered Ca^2+^ signals in the distal compartments (Fig. 1d,c). These Ca^2+^ propagated toward the somatic compartment. Similarly, stimulating all compartments with dopamine triggered Ca^2+^ in all compartments (Fig. 1f,g). In both cases, higher stimulation frequency produced higher frequency response (Fig. 1d,f). With exception for compartments 8 and 9, the Ca^2+^ signals triggered with both glutamate and dopamine have a spike-like form. In the most distal compartments, the stimulation increased the [Ca^2+^] without triggering this spike-like Ca^2+^ signal (Fig. 1e,g).

To distinguish between the model response to glutamate and dopamine, we obtained the input-response frequency (I-f) curve for both stimuli. We varied the input frequency from 0 (no stimulation) to 10 Hz. Only the distal compartments were stimulated with glutamate, while all compartments received the dopaminergic input. The frequency of Ca^2+^ signals in the simplified model in response to glutamate follows an almost linear curve with the input frequency (Fig. 1h, top left panel). In this case, no response was detected in the somatic and proximal compartments. In contrast, for the dopaminergic stimulation, the I-f curve is closer to an all-or-none response that saturates for higher input frequency (Fig. 1h, top right panel). Adding a glutamatergic input with 1 Hz to the distal compartments had a minor effect over the response frequency to the dopaminergic stimulation (Fig. 1h, bottom left panel). A constant dopaminergic input with 1 Hz also increased the response frequency to glutamate without changing the linear relationship (Fig. 1h, bottom right panel). This test shows that while the astrocyte model almost respond linearly to the glutamatergic stimulation, it shows all-or-none response to dopamine. This suggests that astrocytes encode synaptic input in a linear fashion, while encode information conveyed by neuromodulators as an all-or-none. In the presence of neuromodulators, such as dopamine, the response to synaptic input is enhanced.

To understand the differences detected in the Ca^2+^ signals from different compartments, we analyzed the system phase space and compared the dynamics among compartments. The derivation and description of the dynamical system can be found in the Dynamical System Section 3.2. In the case for dynamical system analysis, we set *D*_*c*_ = *D*_*i*_ = 0, that is, without diffusion. We plotted the system nullclines and calculated the equilibrium points considering the disconnected model. The trajectories, however, are simulated with the connected model. The *i*-nullcline is bell-shaped and is the same among the compartments (Fig. 1i). This is consistent with the independence of IP_3_ synthesis to compartment parameters (equation (15)). In contrast, the *c*-nullcline has a plateau form in thicker compartments while it has a sharp increase for thinner compartments (Fig. 1i). Without stimulation, the equilibrium point is stable. Adding either glutamate or dopamine move the *i*-nullcline upwards (Fig. 1i). In both cases, the stimulation changes the type of equilibrium point. In compartment 7, the equilibrium point turns into an unstable focus, and in compartments 8 and 9 they become stable focus points (Fig. 1i). Dopaminergic stimulation has a weaker effect over the *i*-nullcline compared to the glutamatergic stimulation (Fig. 1i). This analysis shows that compartments encode information in distinct ways. While proximal or thicker compartments trigger Ca^2+^ signals with a spike-like shape, distal compartments accumulates the input without amplification, as evidenced by the distinct Ca^2+^ dynamics between compartments. Distal compartments could act then as a source of IP_3_ that can diffuse toward thicker regions. Since dopamine has a weaker effect, dopaminergic stimulation has a modulatory role over the synaptic glutamatergic input, corroborated by the different f-I curves obtained for both neurotransmitters.

### 2.2 Types of Ca^2+^ Signals

Ca^2+^ signals with distinct characteristics and patterns are detected in astrocytes from different brain regions [27–30]. Tehari *et al*. [28] with both experimental data and computational simulations classified Ca^2+^ signals intro four categories. Ca^2+^ responses longer than 70 s was classified as long-lasting responses. Signals with only one peak were classified as single-peak. If the response had several close peaks, they were called multi-peak. Finally, responses with a peak followed by a long-duration plateau were classified as plateau response. Although the computational model used by Tehari *et al*. [28] associated the different types of responses to the dynamics of IP_3_, it is not clear how different IP_3_ dynamics emerge from the same inputs. We explore this question in the present section. We analyzed how changing the parameters that control the intracellular Ca^2+^ and IP_3_ dynamics and the transmembrane current could change the amplitude, duration and ISI of the Ca^2+^ signals in response to glutamatergic input. So, we first studied how the Ca^2+^ efflux rate from the ER through IP_3_ channel receptors (*r*_*C*_, equation (11)) and SERCA pump rate (*v*_ER_, equation (13)) could influence the Ca^2+^ signals characteristics (Section 2.2.1). Next, we explored the IP_3_ degradation rate (*r*_5P_, equation (20)) and the strength of Ca^2+^ inhibition to the IP_3_ synthesized by receptors activation (*K*_*p*_, equations (16) and (17), Section 2.2.2). Third, we tested the influence of NCX current parameters (*α*_NCX_ and *β*_NCX_, equation (2), Section 2.2.3). Finally, we analyzed how the glutamate, dopamine and astrocyte morphology could interact and impact the response type detected (Section 2.2.4).

In the test of this section, we stimulated the distal compartments (7, 8 and 9) of the connected model with glutamate with frequency of 5 Hz applied during the entire simulation period (200 s). Since compartment 7 is representative of the response detected in the intermediate, proximal, and somatic compartments (Extended Data Fig.), we focused here on describing the results from the distal compartments.

#### 2.2.1 Variation of Ca^2+^ efflux rate from ER and SERCA pump uptake rate

In the first test, we modified the Ca^2+^ efflux rate from the ER (*r*_*C*_, equation (11)) and SERCA pump uptake rate (*v*_ER_, equation (13)). With exception for the most distal compartment (9), alteration in the Ca^2+^ uptake rate (*v*_ER_) had just a minor effect on the average amplitude of Ca^2+^ signals (Fig. 2a). In contrast, *r*_*C*_ controls the average amplitude, duration and ISI (Fig. 2a). Below a threshold of *r*_*C*_ = 4.076 *s*^−1^ no Ca^2+^ signal is triggered (Fig. 2a). In compartment 9, the average amplitude, in the case in which a Ca^2+^ is triggered, remains constant (Fig. 2a) for different *r*_*C*_. The same is observed regarding the influence of *r*_*C*_ on the average duration and ISI, however, in this compartment, the parameter *v*_*ER*_ changes the *r*_*C*_ threshold for triggering Ca^2+^ signals (Fig. 2a). This result shows that both Ca^2+^ efflux from ER through IP_3_ receptors and the reuptake by SERCA control the amplitude, duration and ISI of the Ca^2+^ signals.

To confirm the influence of these parameters on the Ca^2+^ signal characteristics, we simulated the astrocyte model taking combinations of *r*_*C*_ and *v*_ER_ (Fig. 2d,e,f). We applied at the distal compartments a glutamatergic input with frequency of 5 Hz. This stimulation triggered Ca^2+^ signals in the three parameter combinations (Fig. 2d,e,f). The frequency of Ca^2+^ signals increased as both parameters also increased, consistent with the average ISI decrease with *r*_*C*_ (Fig. 2c). In addition, the duration of Ca^2+^ also decreased with parameters (Fig. 2d,e,f, right panels). So, increasing both *r*_*C*_ and *v*_ER_ facilitated the generation of Ca^2+^ spikes. Consistent with this, analyzing the phase space, we see that increasing both parameters attenuates the sharp increase detected in the *c*-nullcline of distal compartments (Fig. 2g). That is, the distal compartments dynamics come closer to the dynamics of the intermediate and proximal compartments, facilitating their response. As described in the detailed astrocyte model, this changes moves the distal compartments from the ‘passive’ dynamics to a more ‘active’ one [26].

**Fig. 2.**
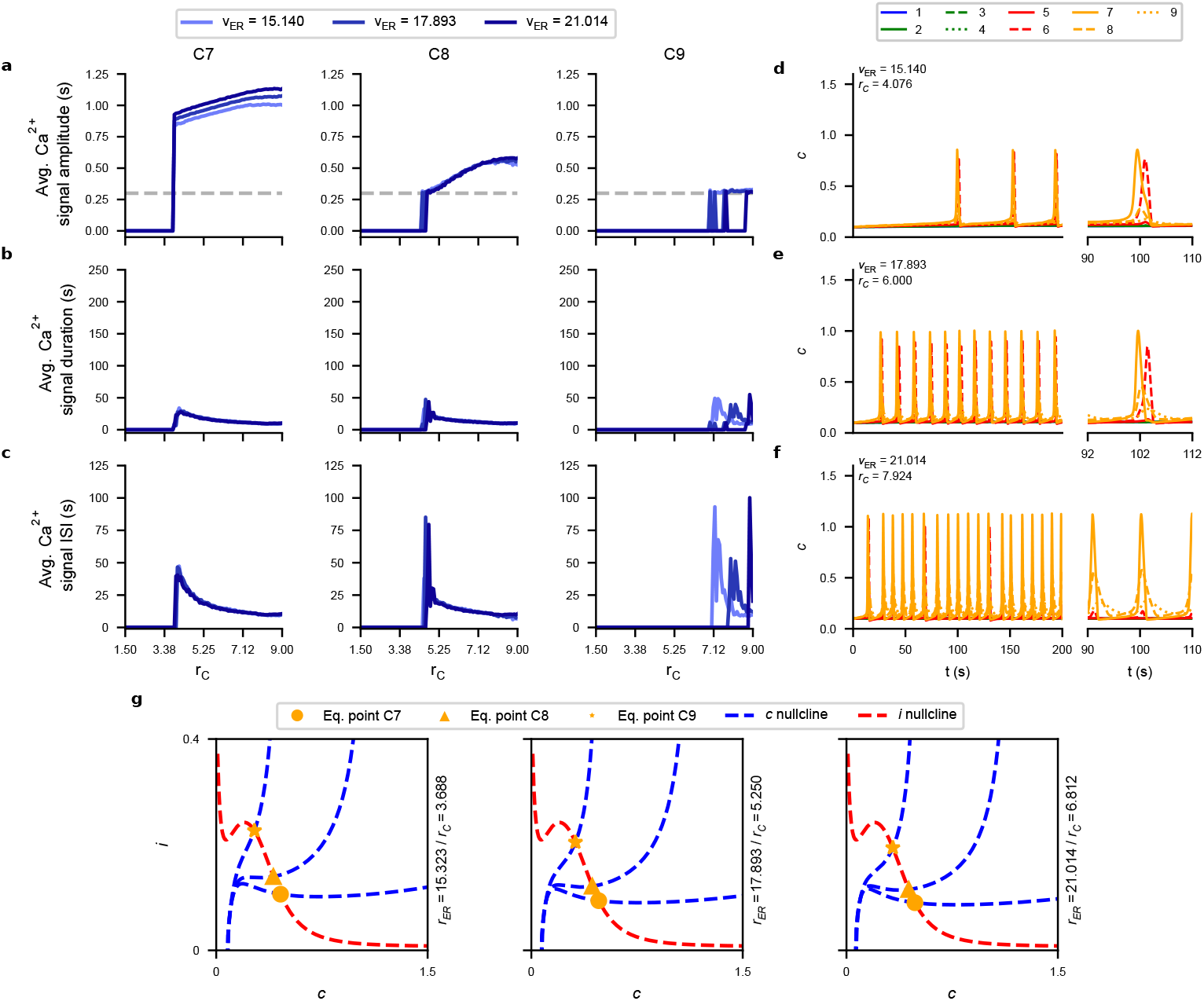
Ca^2+^ efflux rate from ER and SERCA pump uptake rate control the amplitude, duration and ISI of Ca^2+^ signals. **a** Ca^2+^ amplitude, **b** duration and **c** for different values of *v*_ER_ and *r*_*C*_ with glutamatergic stimulation with 5 Hz. **d, c, f** trace of intracellular [Ca^2+^] for three combinations of *v*_ER_ and *r*_*C*_ with glutamatergic stimulation with 5 Hz. Panel on the right show an amplification of a Ca^2+^ **g** Phase space for the same *v*_ER_ and *r*_*C*_ combinations as **d, c, f** with constant glutamatergic input (Extended Data Fig 1).

#### 2.2.2 Variation of IP_3_ degradation rate and the strength of Ca^2+^ inhibition to the IP_3_ synthesized by receptors activation

To test the effect of altering the parameters of IP_3_ dynamics over the Ca^2+^ signals, we varied the IP_3_ degradation rate (*r*_5P_, equation (20)) and the strength of Ca^2+^ inhibition to the IP_3_ synthesized by receptors activation (*K*_*p*_, equations (16) and (17)). Altering the parameters *K*_*p*_ did not impact the Ca^2+^ signal characteristics analyzed (Fig. 3a,b,c). In contrast, increasing the degradation rate *r*_5*P*_, decreased the average Ca^2+^ amplitude, but slightly increased the duration and ISI (Fig. 3a,b,c). However, in any case, setting the highest IP_3_ degradation rate abolished the Ca^2+^ signaling.

**Fig. 3.**
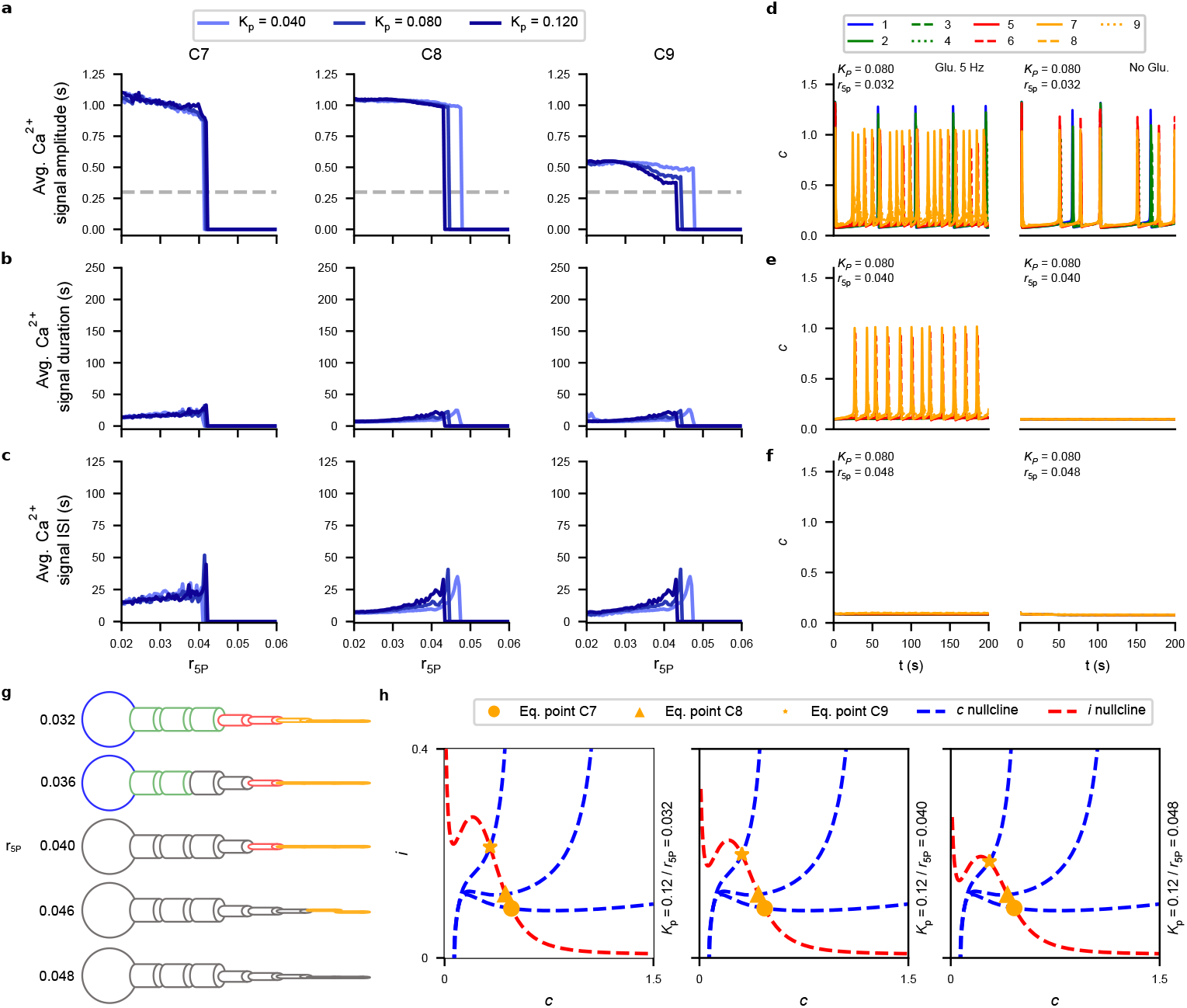
IP_3_ degradation rate controls the astrocytic spontaneous and global responses. The degradation rate *r*_5P_ and strength of Ca^2+^ inhibition of IP_3_ synthesis *K*_*p*_ have a minor effect over the average **a** amplitude, **b** duration and **c** ISI of Ca^2+^ signals. Traces of *c* (representing intracellular [Ca^2+^] for three combinations of *r*_5P_ and *K*_*p*_ with (left) and without (right) glutamatergic stimulation. Lower degradation rates trigger global and spontaneous responses. **g** Global responses are triggered for lower degradation rates (*r*_5P_ = 0.032 *s*^−1^ or *r*_5P_ = 0.036 *s*^−1^). Gray compartments represents no response during simulation. The compartments are color coded as in Fig. 1. **h** Phase space with the same three parameter combinations as **d, e** and **f**.

To explore these results, we simulated the astrocyte model applying glutamate with 5 Hz at the distal compartments. Decreasing both degradation rate and Ca^2+^ inhibition over IP_3_ synthesis from the default values facilitated the response in the entire astrocyte. Ca^2+^ signals were triggered in all compartments, including the soma, with higher activity frequency in the compartments under stimulation and those neighboring them (Fig. 3d). Using parameters closer to the default value (Fig. 3e), the response in the somatic and proximal compartments is abolished, glutamate triggered Ca^2+^ signals only in the distal compartments. Finally, taking the highest values for *r*_5P_ and *K*_*p*_ abolished all the astrocyte response (Fig. 3f). Even without glutamatergic stimulation, we detected Ca^2+^ signals with the first parameter combination (Fig. 3d). So, with this combination the model triggered spontaneous global responses. Glutamatergic input also increased the frequency of response in the proximal, intermediate and somatic compartments (Fig. 3d). Interestingly, disabling the diffusion from the proximal to the distal compartments, but keeping the diffusion form the distal compartments to the proximal ones, decreased the responses in the distal compartments (Extended Data Fig.). This suggests that the spontaneous global responses facilitate the generation of Ca^2+^ signals in the distal compartments. The spontaneous global responses were triggered only with lower degradation rates *r*_5P_ (Fig. 3g). Interestingly, some responses were confined to the somatic and proximal compartments (Fig. 3g, *r*_5P_ = 0.036 *s*^−1^). So, although not affecting the average amplitude, duration and ISI of the Ca^2+^ signals triggered, the IP_3_ dynamics also affect the generation of spontaneous and global responses.

Analyzing the phase space, we detected that changing the IP_3_ parameters altered only the *i*-nullcline (Fig. 3h). Consistent with their effect over the Ca^2+^ signaling, increasing the values of *r*_5P_ and *K*_*p*_ reduced the glutamate effect on moving upwards the *i*-nullcline, so reducing the strength of glutamate in triggering Ca^2+^ signals. For the lowest *r*_5P_, the equilibrium point without stimulation was moved to the regions in which Ca^2+^ responses are triggered, which explained the spontaneous response (Extended Data Fig.).

#### 2.2.3 NCX Current Parameters

Next, we analyzed how changing the parameters of the NCX Ca^2+^ current through the astrocyte membrane would impact the characteristics of the Ca^2+^ signals (equation (2)). In this test the distal compartments were also stimulated with glutamate with frequency of 5 Hz. In compartments 7 and 8, *α*_NCX_ slightly changes the average Ca^2+^ signals amplitude and duration (Fig. 4a,b). *α*_NCX_ had a U relationship with the average Ca^2+^ signals ISI (Fig. 4a). *β*_NCX_ had a minor effect over the Ca^2+^ characteristics in these two compartments. In contrast, in compartment 9, the average Ca^2+^ signal amplitude decreased with *α*_NCX_ and, past a threshold determined by *β*_NCX_, the Ca^2+^ signals were abolished in that compartment (Fig. 4a). Low *α*_NCX_ affected the basal [Ca^2+^] in compartment 9 (altered the equilibrium value of *c*), creating a long-lasting plateau (Fig. 4b). Although not largely affecting the ISI in this compartment, *α*_NCX_ created a peak in the average Ca^2+^ ISI in compartment 9 (Fig. 4c). With exception for compartment 7, there is a threshold *α*_NCX_ value above which the Ca^2+^ signals are abolished. The threshold value of *α*_NCX_ depends on the *β*_NCX_. So, although influencing the amplitude and duration in the most distal compartments, the NCX controls more strongly the timing of the Ca^2+^ signals.

**Fig. 4.**
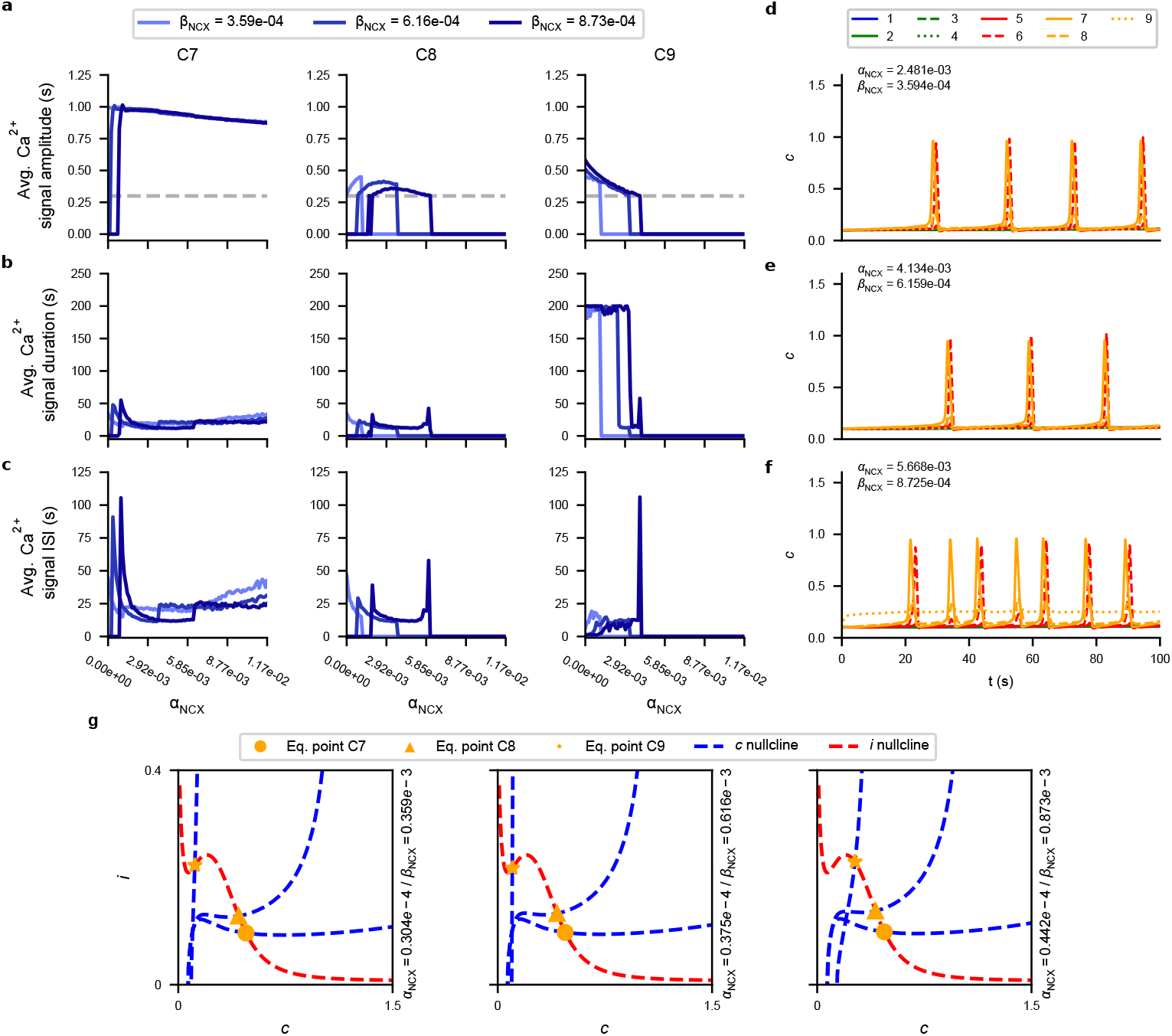
Transmembrane Ca^2+^ current controls the average ISI between Ca^2+^ signals and the response in the distal compartments. The parameters *α*_NCX_ and *β*_NCX_ had different effects between compartments over the average Ca^2+^ **a** amplitude, **b** duration and **c** ISI. **d**,**e**,**f** Traces of *c* (representing intracellular [Ca^2+^]) for three combinations of *α*_NCX_ and *β*_NCX_. **g** Phase space of the three parameter combinations in **d**,**e**,**f**.

Consistent with the U shape observed for the Ca^2+^ signal characteristics, simulating the model applying a glutamatergic input with 5 Hz at the distal compartments produced a similar response. Reducing the values of *α*_NCX_ and *β*_NCX_ increased the frequency of Ca^2+^ signals detected compared to the default values (Fig. 4d,e). Similarly, increasing both *α*_NCX_ and *β*_NCX_ also increased the frequency of Ca^2+^ signals detected (Fig. 4e,f). However, in this case, the intracellular Ca^2+^ concentration (variable *c*) reached a higher steady state in compartment 9, consistent with their effect over compartment 9 basal Ca^2+^ concentration. In the phase space, only increasing both *α*_NCX_ and *β*_NCX_ produced a noticeable difference in the model dynamics, moving the *c*-nullcline of compartment 9 rightwards (Fig. 4g), explaining the increased baseline Ca^2+^ concentration in this compartment.

Taking together, these results show that the three factors analyzed here control different characteristics of the Ca^2+^ responses in astrocyte. While the ER Ca^2+^ mechanism controlled the Ca^2+^ signals amplitude in the more proximal compartments, the IP_3_ degradation influenced the generation of global and spontaneous responses, and the transmembrane current affected the response of the distal compartments and the average Ca^2+^ ISI.

#### 2.2.4 Influence of neurotransmitters and morphology over the Ca^2+^ response type

In the previous sections we analyzed how the mechanisms related to the release and uptake of Ca^2+^ from/to ER, the IP_3_ synthesis and degradation and the transmembrane currents could affect the amplitude, duration, ISI and the location of Ca^2+^ signals. However, in these tests, the astrocyte was stimulate with glutamate. But, as observed in a previous study of our group [26], dopamine triggered global responses and the morphological structure of astrocytes also influenced their response. So, to understand how the global response triggered by dopamine and the astrocyte morphology could be associated with the type of Ca^2+^ signals, we tested whether stimulating the astrocyte with both glutamate and dopamine and disconnecting the astrocyte model would modify the type of Ca^2+^ signal triggered.

We first studied the model response to both glutamate and dopamine. In this test, the entire astrocyte was stimulated with dopamine with frequencies of 5 Hz or 10 Hz and the distal compartments with glutamate also with frequencies of 5 Hz and 10 Hz. As controls, we simulated conditions with only dopamine or only glutamate. To increase the duration of the Ca^2+^ response, we changed the values of the ER parameters to *r*_*C*_ = 4.076 *s*^−1^ and *v*_ER_ = 15.140 *s*^−1^, combination correspondent to the peak of Ca^2+^ signal duration obtained in the test varying *r*_*C*_ and *v*_ER_ (Fig. 2b). Without this modification, we detected no difference in the type of response triggered by the dopaminergic and glutamatergic stimulation (Extended Data Fig.). Glutamate or dopamine alone triggered only single peaks (Fig. 5a), with dopamine triggering global responses and glutamate only in the distal compartments (Fig. 5a). This response is consistent with the patter of release of dopamine and glutamate [34, 36, 37]. With exception for the case with glutamatergic stimulation with 10 Hz, the combination of glutamate and dopamine triggered a plateau response in the distal compartments (Fig. 5a). For the glutamatergic input with 10 Hz and dopamine with 5 Hz, some of the Ca^2+^ signals are double peaks (Fig. 5a). Since the glutamatergic stimulation alone is not enough to produce neither the plateau responses, the dopaminergic input is needed for the this type of response.

**Fig. 5.**
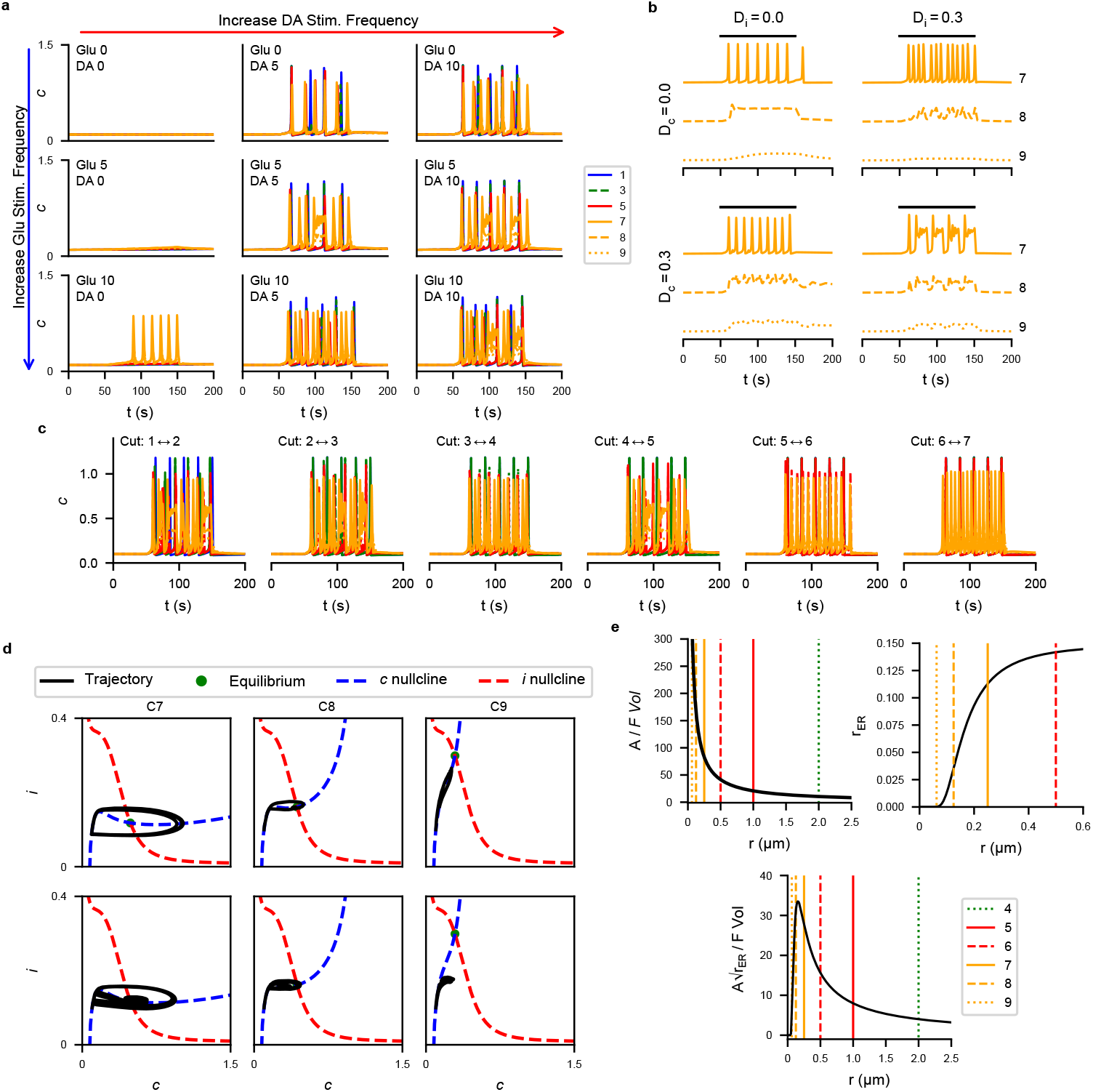
Neurotransmitter type and astrocyte morphology influence the type of Ca^2+^ signal. **a** traces of the intracellular Ca^2+^ concentration (variable *c* with different frequencies of glutamatergic and/or dopaminergic stimulation. **b** astrocyte response to a glutamatergic and dopaminergic inputs both with frequency of 10 Hz with and without Ca^2+^ and/or IP_3_ diffusion. **c** Same test as **b** but blocking diffusion only at specific compartment connections. **d** phase space and trajectory with-out (upper panels) and with (lower panels) diffusion. **e** factors scaling the currents through ER or astrocyte membrane as functions of the compartment radius.

Since different compartments have distinct dynamics (Fig. 1 and Extended Data Fig.) and dopamine trigger global response and is needed for the plateau responses (Fig. 5a), the communication between compartments could be responsible for the type of Ca^2+^ signals detected. To understand how the connection between compartments could influence the Ca^2+^ response, we analyzed the disconnected model stimulating the astrocyte with dopamine and glutamate, both with frequency of 10 Hz with or without Ca^2+^ or IP_3_ diffusion. Without either Ca^2+^ or IP_3_, we observed just single peaks in compartment 7, a single plateau in compartment 8, distinct from the plateau response, and a small fluctuation in compartment 9 (Fig. 5b). Interestingly, with diffusion of only IP_3_, the Ca^2+^ signals were triggered in close proximity, similar to a multipeak responses [28] (Fig. 5b). This suggests that IP_3_ diffusion is more important for the type of Ca^2+^ signal triggered [28]. In addition, disconnecting the distal compartments from the rest of astrocyte had a similar effect as the entire disconnect model (Fig. 5c). Similarly, preventing diffusion from the distal compartments also abolished the plateau response (Fig. 5c). Preventing only Ca^2+^ diffusion did not abolished the plateau response in the distal compartments (Extended Data Fig.). So, the responses triggered in the more proximal regions and the communication between astrocyte regions are needed for the astrocyte to trigger different types of Ca^2+^ signals.

The diffusion effect over the type of Ca^2+^ signal can be interpreted considering the system dynamics (Fig. 5d). Without diffusion, the trajectory of each compartment follows the expected trajectory considering the type of the equilibrium points (unstable focus in compartments 7, stable focus in compartments 8 and 9; Fig. 5d). Connecting the compartments, this trajectory is deviated. In compartment 7, for example, the trajectory is pushed toward the trajectory of compartment 8, creating plateaus in the Ca^2+^ concentration of compartment 7. However, since the equilibrium point of the compartment 7 dynamics is still an unstable focus, its trajectory is again pushed toward the limit cycle around the equilibrium point, terminating the plateau response.

Finally, the behavior difference between compartments can be explained considering the morphological parameters and current scaling factors that control the contribution of each Ca^2+^ mechanisms (ER-or transporter dependent, Fig. 5e). The factor *A/F* 𝒱 (equation (9)) that corrects the NCX current diverges for compartment thinner than compartment 7 (Fig. 1a and 5d). Similarly, the ER-cytosol volume ratio *r*_ER_ is similar for compartments thicker than compartment 7. So, the factor that corrects the Ca^2+^ ER current 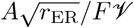 (equation (9)) has a peak between the radii of compartment 7 and 8 (Fig. 5d). So, compartments thinner than compartment 7 will show a greater NCX current compared to thicker compartments while a smaller ER Ca^2+^ current.

Here we present a simplified bidimensional astrocyte model for the simulation of the Ca^2+^ and IP_3_ dynamics. The model Ca^2+^ response replicates the experimental data and previous models [26]. With this model, we simulated Ca^2+^ signals with different characteristics by changing the parameters of Ca^2+^ and IP_3_ dynamics, and the transmembrane current. Interestingly, these factors affected different characteristics of the Ca^2+^ signals, as the amplitude, ISI and location of responses. To generate different types of Ca^2+^ signals, we observed that the global response triggered by dopamine is needed to generate the plateau response. Finally, we propose a mechanism that could explain how the astroyctes control the type o gliotransmitter released.

After the simplification of a biophysical detailed astrocyte model [26], the model is reduced to a two variable system, describing the intracellular Ca^2+^ and IP_3_ concentrations. The simplified model retains both ER- and NCX-dependent mechanisms that generate Ca^2+^ signals in astrocytes. The characteristics of the Ca^2+^ obtained here are consistent with experimental data [16, 28] and with previous computational models [15, 24, 26, 28]. As we observed in a previous study [26], dopamine and glutamate have distinct roles over the astrocyte response. While glutamate released by synapses ensheathed by an astrocyte controls locally the Ca^2+^ signaling [10, 34, 36, 37], neuromodulators could trigger global responses, and create a broad response in the cell [32, 38]. The frequency of glutamatergic input also creates an almost linear relationship with the astrocyte response frequency. In contrast, dopaminergic transmission produces an all-or-none response. In addition, distal compartments act as passive integrators [26] that does not amplify the input by triggering Ca^2+^ signals. Taking together, glutamate could signal to astrocytes the frequency of synaptic activity and be associated with local astrocytic computation and synaptic modulation. In contrast, neuromodulators, such as the dopamine, can encode for contextual information [14, 26], triggering the global responses in astrocytes and, then, could modulate several astrocytic terminals simultaneously.

Changing the parameters controlling the Ca^2+^ (Section 2.2.1) and IP_3_ (Section 2.2.2) dynamics, and the NCX current (Section 2.2.3) differentially altered the characteristics of the Ca^2+^ signals triggered. While the Ca^2+^ dynamics parameters influenced the Ca^2+^ signals amplitude and ISI, the parameters of the IP_3_ dynamics controlled the generation of spontaneous and global responses. Interestingly, the NCX current parameters modified the ISI of Ca^2+^ signals in a U-shape trend, and also the basal Ca^2+^ concentration in the distal region of the astrocyte model. By changing the NCX current parameters, we could interpret this current as an effective Ca^2+^ transport across the membrane, not just a current through the NCX. So, the term corresponding to the NCX (equation (9)) could be modified to represent other transmembrane current present in the astrocyte. So, altering the parameters and matching the Ca^2+^ characteristics, we can use the present model to replicate the response of astrocytes from different brain regions [19–21]. Since the hippocampal astrocytes show a higher frequency of spontaneous responses [19], it could be replicated by changing the parameters associated with IP_3_ dynamics [15]. Similarly, since the basal Ca^2+^ concentration of astrocyte in striatum depends more strongly on the transmembrane Ca^2+^ currents [19], this could be simulated with the present model by changing the parameters of the NCX current, now interpreted as an effective current through the membrane. Although not explored in detail here, the present model offers the opportunity to understand the origin of the Ca^2+^ response diversity and to investigate the role of these diversity in neural systems. As the Izhikevich model for neurons [39], the present astrocyte model could be used to simulate and implement different types of astrocyte in neural networks and investigate their role on the regulation and mediation of cognitive functions. However, the computational models present in the literature simulate a general abstract astrocyte [15, 24, 26, 32], disregarding the diversity of responses described in the astrocyte literature [19–21]. Future works of our group will address this question.

In addition to the response diversity, astrocytes also show different patterns of Ca^2+^ signals [27–30]. As described by Taheri *et al*. [28], the Ca^2+^ signals can be classified into single peaks, multipeaks, plateau and long-lasting responses. With a computational model, they showed that different IP_3_ time courses are associated with these response types. However, it is not clear how the same inputs could determine the IP_3_ time course and change the response type. Here we show that with a neuromodulator that triggers global responses, as the dopamine in the present model, coupled with the glutamatergic input, different types of Ca^2+^ signals emerge. Stimulating the model with dopamine and glutamate triggered responses similar to the plateau response [28]. Although the different types of Ca^2+^ signals are also detected in the somatic and proximal regions [27–30], plateau responses were detected only in distal compartments. Since these are the compartments receiving directly glutamatergic input, it could explain why only these compartments triggered other types of response. As shown before, the astrocytic morphology can change the compartment response [26]. Therefore, using more complex morphologies with more branches could induce other types of response in the somatic and proximal compartments. So, to understand the Ca^2+^ signaling in astrocytes, it is essential to account for the cell morphology.

Since blocking the diffusion of Ca^2+^ and IP_3_ or just disconnecting the distal compartments abolished the plateau response, the global Ca^2+^ signals seems to be necessary for different patterns of Ca^2+^ responses. Interestingly, blocking only IP_3_ was enough to prevent the generation of the plateau response. So, although the Ca^2+^ concentration is the most studied component in the astrocyte excitability [10], the present result suggests that the IP_3_ should be considered as one of the ‘computational’ units in astrocytes. This perspective is consistent the attenuation of Ca^2+^ response with IP_3_ buffering or knockout of its receptor [22, 40]. This also explains why the global responses triggered in astrocyte seems to be disconnected and encompass local simultaneous but large scale responses [16]. The relationship between global responses triggered by a neuromodulator and the type of Ca^2+^ signal detected is a prediction of our model that requires experimental test.

The model present here can be used to study astrocyte excitability, showing a good approximation to the Ca^2+^ response in astrocyte seem in other computational models and experimental data. Since the model has only two variables, it enables the use analytical tools to study its behavior. With the simplified model it is possible to study more readily the parameter space and match the model response to the diversity of astrocyte Ca^2+^ signaling observed experimentally. Here we present a hypothesis linking the type of Ca^2+^ signal with the global response and neuromodulators.

## 3 Model Description

The original biophysically detailed compartmental astrocyte model is described elsewhere [26]. That model is composed of ten state-variables representing the intracellular Ca^2+^ concentration ([Ca^2+^]_i_), extracellular Ca^2+^ concentration ([Ca^2+^]_e_), intra-ER Ca^2+^ concentration ([Ca^2+^]_ER_), intracellular IP_3_ concentration ([IP_3_]), fraction of opened IP_3_ receptors (*h*), intra- and extracellular Na^+^ concentration ([Na^+^]_i_ and [Na^+^]_e_), intra- and extracellular K^+^ concentration ([K^+^]_i_ and [K^+^]_e_) and the astrocyte membrane potential (*V*). Additionally, there are two variables for extracellular glutamate concentration ([Glu]) and dopamine concentration ([DA]). For each of these variable, there is a differential equation describing its rate of change, totaling twelve differential equations. In addition, all compartments are connected by diffusion current densities.

### 3.1 Model Simplification

To derive the simplified model, we first noted that the variables [Na^+^]_i_, [Na^+^]_o_, [K^+^]_i_, [K^+^]_o_, *V*, [Ca^2+^]_o_ are fast compared to the intracellular [Ca^2+^] and [IP_3_] [26], so we substituted them by their mean value (Extended Data Fig. and Supplementary Table). Similarly, comparing the temporal evolution of the variable *h* for different stimuli (Extended Data Fig.), we substituted it by its mean value with a glutamatergic and dopaminergic inputs both with frequency of 5 Hz.

So, the the equation for the current through the Na^+^-Ca^2+^ exchanger (NCX) in the original model [26] was reduced to:

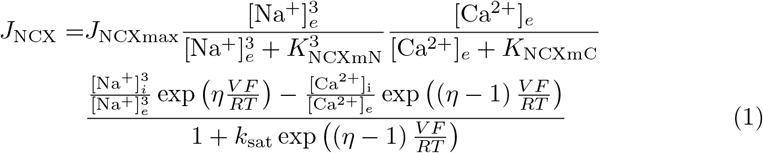

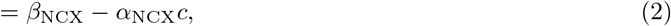

where *α*_NCX_ = 1.169 10^−4^*MA/m*^2^ and *β*_NCX_ = 1.232 10^−5^*MA/m*^2^.

Next, to simplify the [Ca^2+^]_ER_ we first noted that the differential equations describing its time evolution could be rewritten as follows:

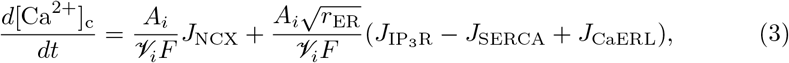

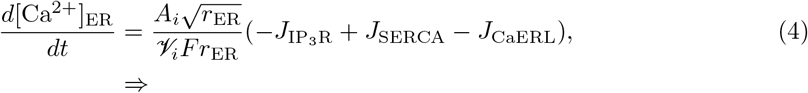

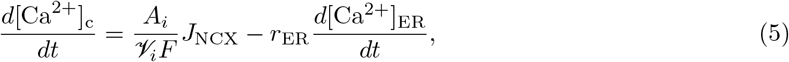

Supposing *J*_NCX_ *≈* 0 and rearranging the terms we get:

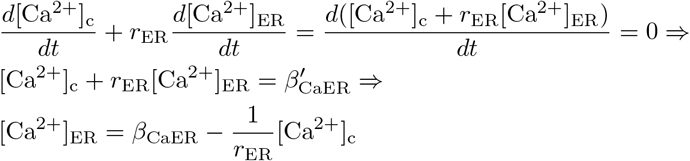

Where 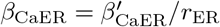. This is a reasonable approximation since the NCX mechanism has a minor effect in the thicker and somtic compartments, so the fluctuations in the intracellular [Ca^2+^] are due to the ER mechanism (Extended Data Fig.). This approximation is worse for the distal compartments (Extended Data Fig.).

However, for a better approximation of [Ca^2+^]_ER_ as a function of [Ca^2+^]_c_, we scaled the *r*_*ER*_ by the factor *α*_CaER_, and imposed a dependence of *α*_CaER_ and *β*_CaER_ with the *r*_*ER*_ as follow:

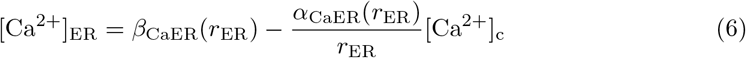

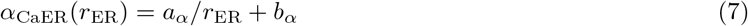

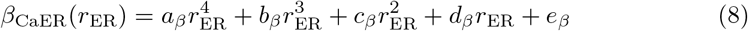

where *a*_*α*_ = − 1.03 10^−4^, *b*_*α*_ = 1.00, *a*_*β*_ = − 3.58 10^5^, *b*_*β*_ = 1.52 10^5^, *c*_*β*_ = −2.19 10^4^, *d*_*β*_ = 1.22 10^3^, and *e*_*β*_ = 1.90 10, after normalization of [Ca^2+^]. These parameters were fitted to reduce the mean squared error calculated between the [Ca^2+^]_ER_ temporal series and the approximation simulating the model with a glutamatergic and dopaminergic input at 5 Hz (Extended Data Fig.).

With the above mentioned modifications, the original biophysically detailed astrocyte model was simplified to a system of two differential equations for the intracellular [Ca^2+^] and [IP_3_]. We also modified the parameters values to restrict the value of [Ca^2+^] and [IP_3_] to around 1 by dividing them by their maximum amplitude in response to a glutamatergic and dopaminergic stimulation with 5 Hz. Since the normalized variables do not have a physical unit, we substituted the [Ca^2+^] by the *c* variable and [IP_3_] by the *i* variable.

So, the time evolution of *c* is described by the following equation:

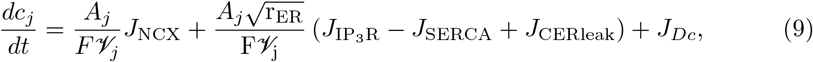

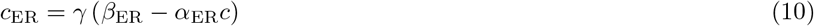

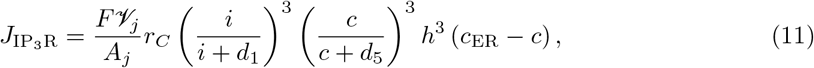

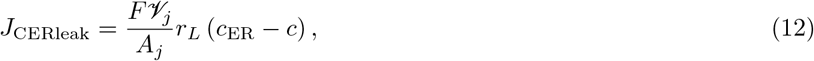

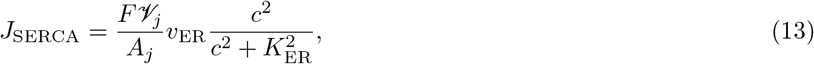

where *A*_*j*_ is the surface area of compartment *j, F* = 96, 500*C/M* the Faraday’s constant, 𝒱_*j*_ the volume of compartment, *J*_NCX_ the current through NCX, *J*_IP_3 R the current through the IP_3_ channels on the ER membrane, *J*_SERCA_ the uptake current by SERCA pump, *J*_CERleak_ the leak current from ER, *r*_*C*_ = 6 *s*^−1^ the Ca^2+^ efflux rate from ER, *d*_1_ = 0.065, *d*_5_ = 0.107, *h* = 0.8, *r*_*L*_ = 0.11 *s*^−1^ it the Ca^2+^ leak current Ca ^2+^ flux rate, *v*_ER_ = 18.782 *s*^−1^ the SERCA uptake rate, *K*_ER_ = 0.116. The factor *γ* is adjusted for each compartment to impose equilibrium at the beginning of the simulation (Supplementary Table).

The *r*_ER_ is defined [24, 26] as:

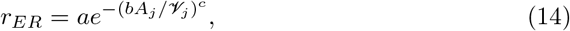

where the parameters *a* = 0.15, *b* = 0.073 µm and *c* = 2.34 were fitted using experimental data [25].

The equation describing the time evolution of *i* is:

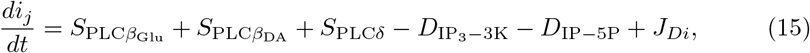

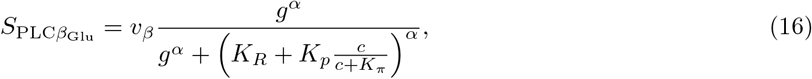

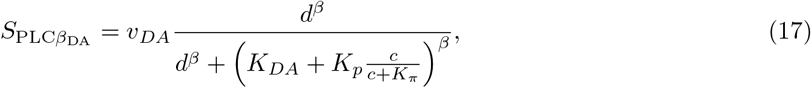

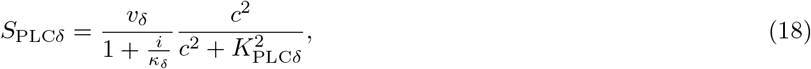

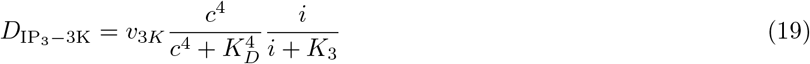

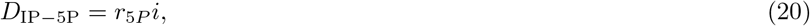

where 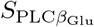 is the synthesis of IP_3_ by the PLC*β* activated by the metabrotropic glutamatergic receptor, *v*_*β*_ = 0.211 *s*^−1^ is the IP_3_ synthesis rate by the mGluR, *α* = 0.7, *K*_*R*_ = 0.104 *·* 10^−2^, *K*_*π*_ = 0.8214, 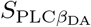 is the synthesis of IP_3_ by the PLC*β* activated by the dopaminergic receptor (D_1_ or noradrenergic *α*_1_), *v*_*DA*_ = 0.013 *s*^−1^, *β* = 0.5, *K*_*DA*_ = 5 *·* 10^−3^, *S*_PLC*δ*_ is the synthesis of IP_3_ by the PLC*δ, v*_*δ*_ = 0.013 *s*^−1^ is the IP_3_ synthesis rate by PLC*δ, κ*_*δ*_ = 0.782, *K*_*P LCδ*_ = 0.137, *D*_IP_3 −3K is the degradation of IP_3_ by the enzyme IP_3_-3K, *v*_3*K*_ = 1.043 *s*^−1^ is the degradation, *K*_*D*_ = 0.958, *K*_3_ = 0.522, rate *D*_IP−5P_ is the degradation of IP_3_ by the enzyme IP-5P, *r*_5*P*_ = 0.04 *s*^−1^ is the degradation rate, and *J*_*Di*_ is the diffusion of IP_3_ to other compartments.

The extracellular glutamatergic *g* and dopaminergic *d* concentrations are given by:

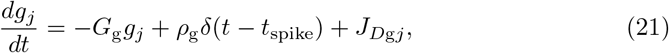

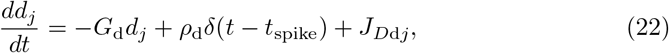

where *g*_*j*_ represents the extracellular glutamate concentration of compartment j, *G*_g_ = 100 *s*^−1^ is the clearance rate of glutamate, *ρ*_g_ = 0.4 *·* 10^−3^ is the amount of glutamate released by each presynaptic spike, *J*_*D*g_ = 4*·*10^−4^*s*^−1^ is the diffusion of glutamate between extracellular compartments, *d*_*j*_ represents the extracellular dopamine concentration of compartment j *G*_d_ = 4.201 *s*^−1^ is the clearance rate of dopamine, *ρ*_d_ = 1 10^−3^ is the amount of dopamine released by each presynaptic spike and *J*_*D*d_ = 13.8 *s*^−1^ is the diffusion of dopamine in the extracellular space. Presynaptic spikes will be simulated as *δ*(*t*− *t*_spike_), in which *t*_spike_ indicates the spike times following a exponential distribution.

The diffusion density currents between compartments are calculated as follows:

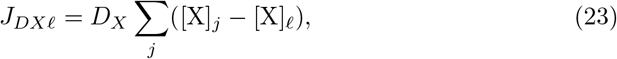

where *𝓁* is the index of the compartment from/to which the ion/molecule X (*c, i, g* or *d*) is diffusing, *j* is the index of the compartments connected to the compartment *𝓁*, and *D* is the diffusion coefficient (in s^-1^) of the ion/molecule X.

A response was classified as a Ca^2+^ signal if it surpassed the threshold *c*_*th*_ = 0.3.

### 3.2 Dynamical System Analysis

By reducing the ten state-variables from the original model to two state-variables in the simplified model, it becomes possible to apply dynamical system tools to study the Ca^2+^ and IP_3_ dynamics for different inputs and morphological parameters.

We investigated the effect of *r*_*ER*_ and glutamatergic and/or dopaminergic over the Ca^2+^ and IP_3_ dynamics with the dynamical system analysis. All algebraic steps (calculation of the nullclines and the system jacobian matrix) are described in the Supplementary Materials. In these tests we disconnected the astrocyte compartments by setting all diffusion parameters to zero. Both glutamatergic and dopaminergic stimuli were calculated and simulated as constant values (Extended Data Fig.). So, as a base condition, we set *g* = *d* = 0.

### 3.3 Computational and Numerical Methods

To calculate the numerical solution of the system of differential equations, we used the 4th-order Runge-Kutta method with time step *dt* = 0.01 *ms*. All numerical routines were implemented in Python 3.7 with the Numpy and Scipy packages. In addition, we used the Numba package with JIT compilation and GPU to speed up the simulations. Graphs were made with the Matplotlib package. All codes are available at GitHub.

## Acknowledgments

T.O.B. is supported by a São Paulo Research Foundation (FAPESP) PhD scholarship (grant 2021/12832-7). A.C.R. is partially supported by a Brazilian National Council for Scientific and Technological Development (CNPq) Research Productivity Grant (# 303359/2022-6). This paper was produced as part of the activities of FAPESP Research, Innovation and Dissemination Center for Neuromathematics (Grant 2013/07699-0).

## Declarations

The authors declare no conflict of interest. Code for model implementation and all analyses is available on GitHub. T.O.B. and A.C.R. conceptualized, led the overall project and wrote the paper.

